# Cryo-electron microscopy structure of the lipid droplet-formation protein seipin

**DOI:** 10.1101/418236

**Authors:** Xuewu Sui, Henning Arlt, Kelly P. Brook, Zon Weng Lai, Frank DiMaio, Debora S. Marks, Maofu Liao, Robert V. Farese, Tobias C. Walther

**Author notes:** These authors contributed equally (Maofu Liao, Robert V. Farese, Tobias C. Walther). Address correspondence to: Maofu Liao, Robert V. Farese Jr. (.), and Tobias C. Walther.

## Abstract

Sui *et al*. report the cryo-EM structure of the conserved luminal domain of the lipid droplet (LD)-formation protein seipin. The structure reveals key features of this domain and suggest a new model for seipin’s role in LD formation.

Metabolic energy is stored in cells primarily as triacylglycerols in lipid droplets (LDs), and LD dysregulation leads to metabolic diseases. The formation of monolayer-bound LDs from the endoplasmic reticulum (ER) bilayer is poorly understood, but the ER protein seipin is essential to this process. Here, we report a cryo-electron microscopy structure and functional characterization of *D. melanogaster* seipin. The structure reveals a ring-shaped dodecamer, with the luminal domain of each monomer resolved at ∼4.0 Å. Each luminal domain monomer exhibits two distinctive features: a hydrophobic helix positioned towards the ER bilayer, and a β-sandwich domain that has structural similarity with lipid-binding proteins. This structure, and our functional testing in cells, suggest a model in which seipin oligomers initially detect forming LDs in the ER via hydrophobic helices and subsequently act as membrane anchors to enable lipid transfer and LD growth.

## INTRODUCTION

Nearly all organisms have the capacity to buffer fluctuations in the availability of energy by storing highly reduced carbons in the form of fats, such as triacylglycerol (TG) in lipid droplets (LDs) (Henne et al., 2018; Onal et al., 2017; Walther et al., 2017). Although LDs can be formed in most cells, the structural mechanisms for the biogenesis of this organelle remain largely unknown. In the most widely held model for LD formation (Walther et al., 2017), TGs and other neutral lipids are synthesized by enzymes in the ER membrane and phase-separate to form an oil lens within the ER bilayer. Lenses subsequently grow and bud toward the cytosol, forming a monolayer-bound LD that can be targeted by specific proteins (Henne et al., 2018; Joshi et al., 2017; Walther et al., 2017). ER proteins are thought to be crucial for controlling the LD formation process, but little is known about the underlying mechanisms for generating a relatively homogeneously sized population of LDs.

The ER protein seipin is a central player in LD formation. Seipin, encoded by the lipodystrophy gene *Bernadelli-Seip congenital lipodystrophy type 2* (*BSCL2*) (Gomes et al., 2004; Magre et al., 2001b), is an integral membrane protein with short N-and C-terminal segments in the cytosol, two transmembrane helices, and an evolutionarily conserved ER luminal domain (Fig. 1A) (Lundin et al., 2006). Missense mutations of human *BSCL2/SEIPIN* that lead to lipodystrophy occur mainly in the ER luminal region (e.g., L91P, A212P), suggesting this part of seipin is crucial for its function. In fly and mammalian cells, seipin forms mobile foci in the ER that are recruited to and stabilized at sites of nascent LD formation, where these foci appear to be required for LD growth (Grippa et al., 2015; Holtta-Vuori et al., 2013; Salo et al., 2016; Wang et al., 2014; Wang et al., 2016). In the absence of seipin, cells form many small LDs, possibly due to premature budding, that fail to grow (Grippa et al., 2015; Salo et al., 2016; Wang et al., 2016). Cells also have so-called supersized LDs that likely form from coalescence of the smaller LDs (Fei et al., 2008; Grippa et al., 2015; Salo et al., 2016; Szymanski et al., 2007; Wang et al., 2016; Wolinski et al., 2011). Seipin deficiency also leads to the recruitment of aberrant proteins to LDs and, possibly, to alterations in ER Ca^2+^ homeostasis and lipid metabolism, such as the accumulation of phosphatidic acid (Fei et al., 2011; Han et al., 2015; Wolinski et al., 2015). Previous studies of seipin led to different models for its molecular function, including acting as a regulator of the ER Ca^2+^ pump SERCA, a molecular scaffold and regulator of lipid metabolism enzymes, and a structural protein facilitating LD growth at ER-LD contact sites (Bi et al., 2014; Pagac et al., 2016; Sim et al., 2012; Talukder et al., 2015; Wang et al., 2016). However, despite considerable efforts, the molecular function of seipin in LD biogenesis remains unclear.

**Fig. 1.**
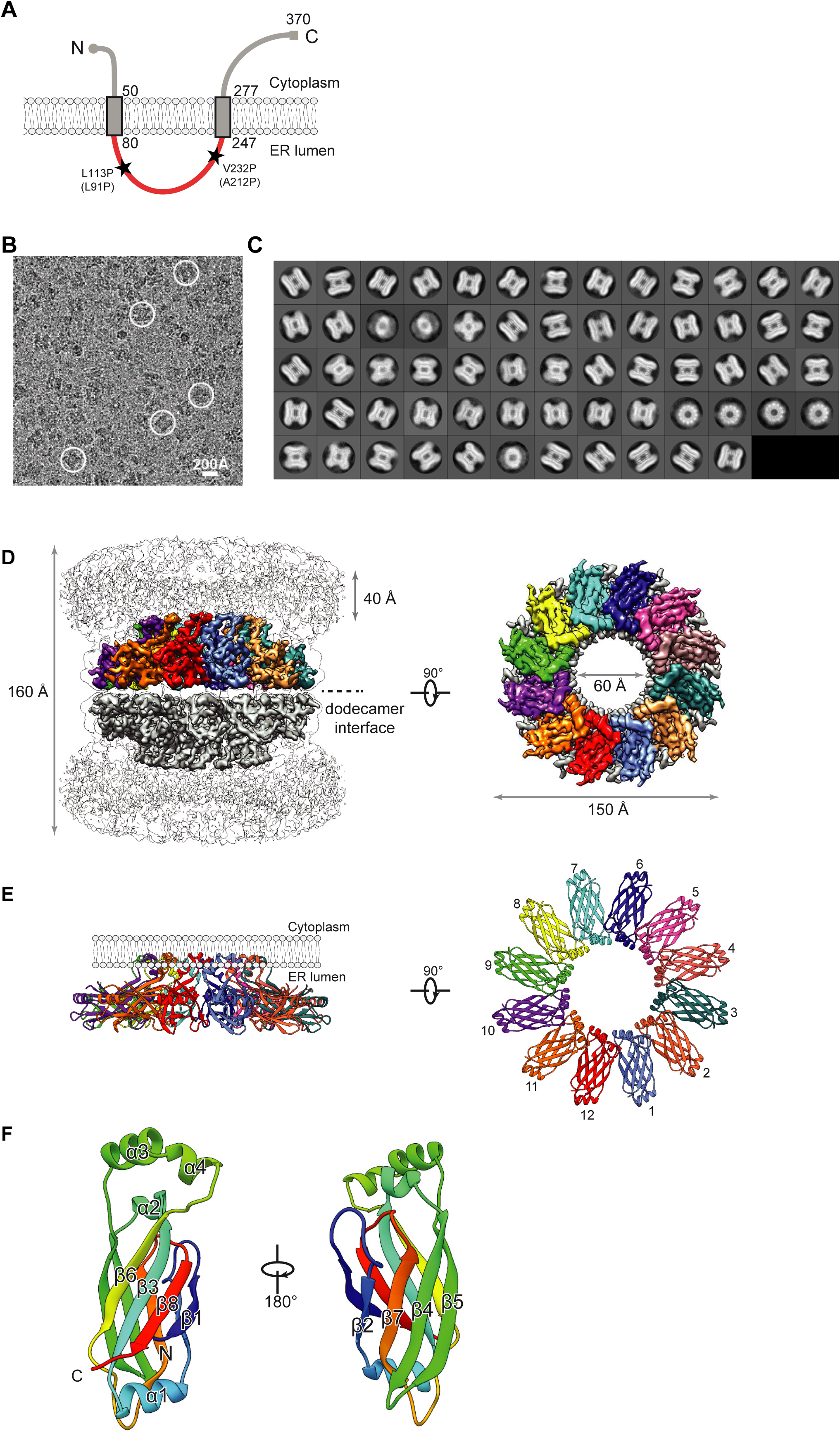
A cryo-electron microscopy map and molecular model of a seipin. **(A)** Model of *D. melanogaster* seipin topology in the ER membrane. The evolutionarily conserved ER-luminal domain is in red. Human pathogenic missense mutations L91P and A212P and their equivalent positions in *Drosophila* seipin are shown. **(B)** Representative cryo-EM image of purified seipin in digitonin. White circles indicate representative particle images of seipin. **(C)** 2D averages calculated with all seipin particles from the combined data sets. The box dimension is 335 Å. **(D)** Unsharpened (transparent) and sharpened (solid colored) cryo-EM density maps of a seipin oligomer. The barrel-like structure is a head-to-head dimer of dodecamers, interacting via the luminal domains. The 40-Å region indicated at the top represents the poorly resolved transmembrane region. Each monomer in the upper dodecamer ring is shown in different colors to assist with visualization. The *en face* view on the right is shown from the perspective of the ER membrane. **(E)** Ribbon diagram side view of the luminal domains positioned towards the ER membrane. To the right is a ribbon diagram top view of the luminal domains. **(F)** Model and structural elements of seipin monomers. The monomer model contains residues 88– 240 of seipin, corresponding to the ER luminal domain. Each luminal domain contains a β-sheet sandwich of eight β-strands and four α-helices. Helices 3 and 4 comprise hydrophobic sequences positioned at the ER luminal leaflet.

As a step toward unraveling seipin’s function, we sought to elucidate its molecular structure. Utilizing cryo-electron microscopy, we report here a structural model of *Drosophila melanogaster* seipin, solved for the luminal domain at ∼4.0 Å resolution. This structure reveals seipin to form a dodecamer that positions multiple hydrophobic helices near the ER bilayer and a β-sandwich folded domain with similarity to lipid-binding proteins. We use several approaches to validate this structure, and we test several of its key features *in vitro* and in cells. Our results suggest a functional model for how seipin functions to detect forming LDs and promote their growth.

## RESULTS

### Determination of a cryo-electron microscopy map of *Drosophila melanogaster* seipin

We purified recombinant *D. melanogaster* seipin in detergents, and gel-filtration chromatography revealed it to be an oligomer (Fig. S1, A and B), similar to what was reported in other species (Binns et al., 2010; Sim et al., 2014). Consistently, negative-stain electron microscopy (EM) analyses showed that seipin particles were monodisperse, and two-dimensional (2D) averages demonstrated distinct views with round or multi-layer barrel shapes (Fig. S1 C).

We next analyzed purified seipin by cryo-electron microscopy (cryo-EM). The 2D averages of cryo-EM particle images (Fig. 1, B and C) appeared similar to those obtained by using negative-stain EM. Three-dimensional (3D) classification of cryo-EM images of seipin purified in digitonin or n-dodecyl β-D maltoside (DDM) detergents showed similar overall conformations, allowing for combined image-processing of these two cryo-EM data sets (Fig. S1 D).

An initial round of 3D classification, performed without symmetry, demonstrated a multi-layer barrel-shaped protein complex containing two middle layers, which correspond to the luminal domains, and top and bottom layers, which are less well resolved and represent the transmembrane (TM) domains. Because 12 repetitive features were clearly discernable in the 2D averages in round-shaped view, we subsequently applied D12 symmetry for further processing (Fig. S1 D). Focusing on the middle layers where the EM density was best resolved, we selected a homogeneous subset of particles to generate a final cryo-EM map at an overall resolution of ∼4.0 Å (Fig. 1 D; Fig. S2 A-C). This map revealed that the barrel-shaped structure consists of two dodecamer rings, interacting head-to-head via seipin luminal domains. This dimer of dodecamers is likely due to non-physiological contacts, inasmuch as we also found single ring dodecamers in protein preparations (data not shown), and it seems unlikely that seipin rings would interact across the lumen of the ER.

The cryo-EM map for most of the seipin luminal domains is of high quality, showing large amino acid side-chain densities and well-separated β-strand density, whereas the loop regions distal from the symmetry axis appear more flexible with lower resolution (Fig. S2 A and D). The N-and C-terminal sequences located on the cytoplasmic side of the ER membrane, as well as the transmembrane domains, were poorly resolved in the density map, likely due to their conformational flexibility.

### A molecular model for *Drosophila melanogaster* seipin

The quality of the cryo-EM map allowed us to build an atomic model of monomeric seipin, spanning amino acid residues 88–240, corresponding to the ER luminal domain. Starting from a manually built, partial model corresponding to the highest resolution region, the combination of Rosetta *de novo* model building (Wang et al., 2015) and model completion with RosettaES (Frenz et al., 2017) confidently placed nearly the entire monomeric sequence. This monomeric model was subsequently refined in the context of the entire symmetric oligomeric assembly (Fig. 1, E and F).

The molecular model of *D. melanogaster* seipin reveals three prominent features. First, the protein forms a ring-shaped dodecamer of laterally interacting subunits, with an overall diameter of ∼150 Å (Fig. 1, D and E). Second, in each seipin monomer, a helical, hydrophobic region (α3 and α4, residues 175–192) of the luminal domain is oriented towards and adjacent to the ER luminal membrane leaflet (Fig. 1 E). Third, the remainder of each luminal domain consists of two β-sheets, each containing four anti-parallel β-strands packed against each other (Fig. 1 F, Fig. S2 D). This region of seipin forms a similar fold as the lipid-binding domain of Niemann-Pick type C2 protein (NPC2; Figure 4F) (Xu et al., 2007). In agreement with this result, *Drosophila* seipin exhibits weak similarities to NPC2 in hidden Markov model searches with the HHPred algorithm (Soding et al., 2005).

To test the structural model, we investigated the interface between monomers. Our model predicts that Tyr230 forms a π-cation interaction with Arg170 and a hydrogen bond with the peptide backbone amino group of Tyr171 (Fig. 2A). To test the requirement of these residues for seipin oligomer formation, we expressed and analyzed mutant forms of seipin by fluorescence-detection size exclusion chromatography (FSEC) of GFP fusion proteins expressed in human cells. Tyr171Ala resulted in a complete shift of the gel filtration peak to a lower molecular weight, indicating weakened multimerization (Fig. 2 B). Expression of either Tyr230Ala or Arg170Ala resulted in a partial multimerization defect. The model also shows Arg165 forming a π–cation interaction with Phe94 on the neighboring monomer (Fig. S3 B). However, Arg165Ala mutation did not alter seipin oligomerization as analyzed by FSEC (Fig. S3 C), and either Arg165Ala or Phe94Ala only showed an effect on oligomerization when combined with a Tyr230Ala mutation, suggesting a less critical role for Arg165 and Phe94 in oligomer formation (Fig. S3 C). Taken together, these results indicate that the seipin structural model correctly predicts key interactions between seipin monomers that contribute to its oligomeric structure *in vitro*.

**Fig. 2.**
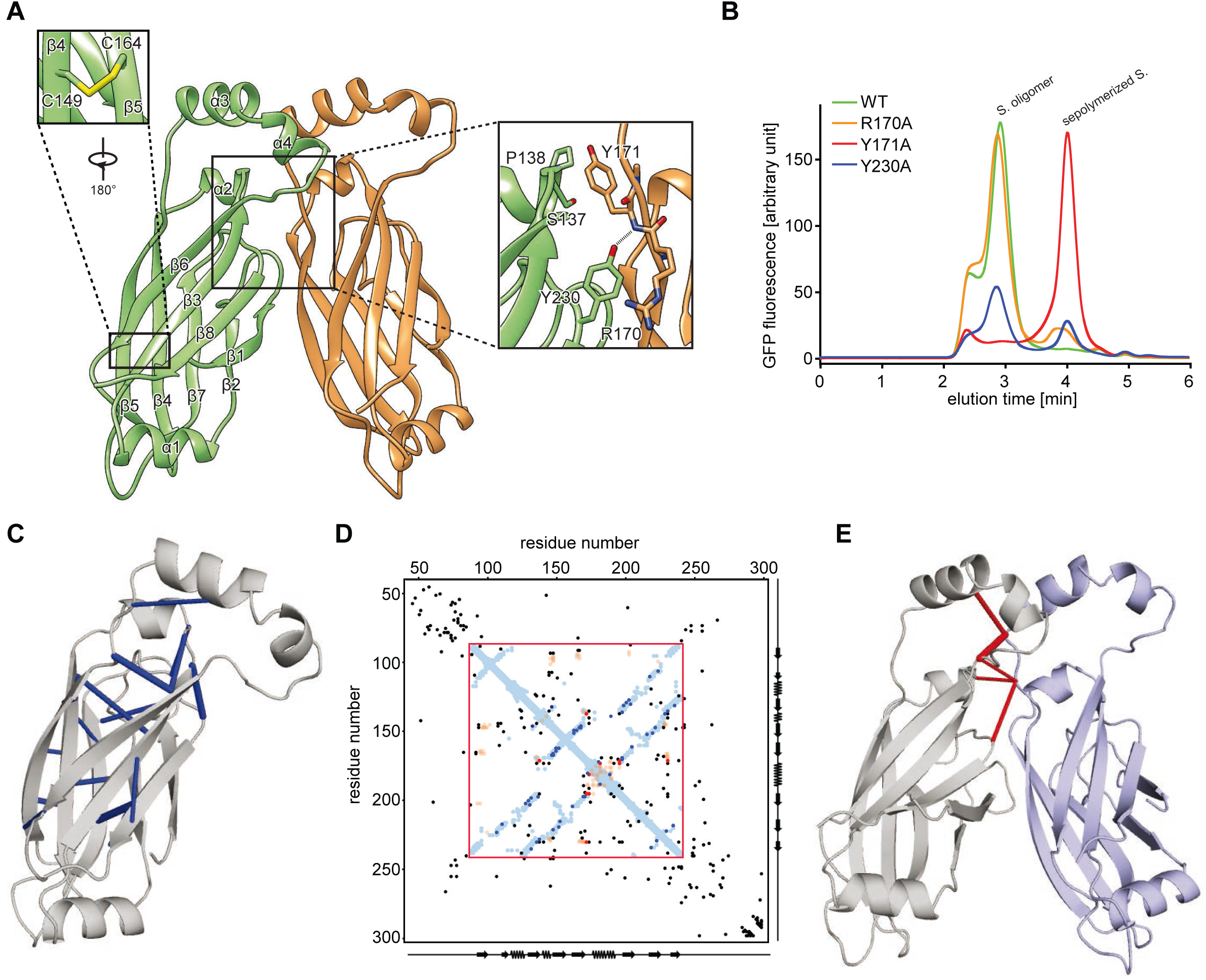
Analysis of seipin oligomer’s inter-and intra-molecular interactions. **(A)** Molecular model of interactions between two seipin monomers. Enlarged views in the boxed regions show the intramolecular disulfide bond (C149 and C164) and key interactions between monomers (discussed in text). **(B)** Fluorescence-based gel-filtration analyses of seipin variants expressed in HEK cells. Mutations (discussed in text) tested include residues at the oligomerization interface shown in **A. (C)** Molecular model of seipin with significant *intramolecular* evolutionary couplings, as revealed by EV fold analyses (Marks et al., 2011; Marks et al., 2012), shown in blue. **(D)** Overlay of seipin’s amino acid evolutionary couplings with distances derived from the molecular model for *D. melanogaster* seipin. Amino acid positions, as well as secondary structure elements, are shown on the x and y axes. Sequence in the red box represents the part of seipin resolved in the cryo-EM structure. Blue-shaded dots represent residues closer than 5Å in the model within a seipin monomer, and orange-shaded dots are residues closer than 5Å between two monomers. Black, blue and red dots represent the top 125 evolutionary couplings, with the most significant ones shown in **A** (blue), and **C** (red). **(E)** Molecular model of seipin with significant *intermolecular* evolutionary couplings shown in red.

As an alternative means to validate the seipin model, we analyzed 1591 seipin protein sequences from different species for evolutionary co-variation that predict physical constraints of protein structure (Marks et al., 2011; Marks et al., 2012). Remarkably, overlaying of the seipin evolutionary couplings with proximities of residues derived from our atomic model showed extensive overlap for residues close within a monomer (Fig. 2 C, D (within the red box) and E], thus supporting the validity of our model. Additionally, the model placed two cysteines (C149 and C164) in close spatial proximity, consistent with density in the cryo-EM map indicating a disulfide bond between these residues (Fig. 2 A, Fig. S2 D). This analysis also was consistent with our findings for important residues interacting at seipin monomer interfaces, as it showed that Tyr230 is evolutionarily highly conserved (Fig. S3 A), and four of the five residues with most enriched evolutionary couplings are in this region, showing for instance a strong coupling between Tyr230 and Tyr171 (Fig. 2 E).

### The hydrophobic helices of the seipin luminal domain bind monolayers of lipid droplets *in vitro* and in cells

The positioning of multiple hydrophobic helices close to the ER bilayer is a distinctive feature of the seipin structure. The hydrophobic helical region of each seipin monomer (Fig. 3 A) exhibited a high degree of evolutionary conservation with respect to hydrophobicity (Fig. 3 B and C). Similar helices with large hydrophobic residues from other LD proteins bind phospholipid packing defects of the LD surface monolayer (Prevost et al., 2018). We thus hypothesized that these hydrophobic helices function to detect packing defects of the ER membrane due to neutral lipid accumulation and lens formation. To test this hypothesis, we incubated a peptide from the seipin hydrophobic helix (seipin-HH) sequence with artificial LDs and found it bound with similar efficiency as a *bona fide* LD-targeting helix (designated P2) derived from the LD protein CCT1 (Prevost et al., 2018) (Fig. 3 D and E). In contrast, a seipin-HH mutant with three negatively charged Asp residues (seipin-HH 3D) failed to bind. We also found that the seipin-HH peptide, but not the seipin-HH3 D mutant, bound predominantly to monolayers of LDs contained in giant unilamellar vesicles (GUVs) (Prevost et al., 2018) (Fig. 3 F and G). Consistent with these results, seipin-HH expressed as a *mCherry* fusion protein targeted to LDs in *Drosophila* S2 cells similar to the M-domain of CCT1 (Kory et al., 2015), whereas seipin-HH 3D did not (Fig. 3 H and I). These results support the hypothesis that the seipin hydrophobic helices serve to recognize and target the protein to phospholipid packing defects of lipid lenses within the ER bilayer.

**Fig. 3.**
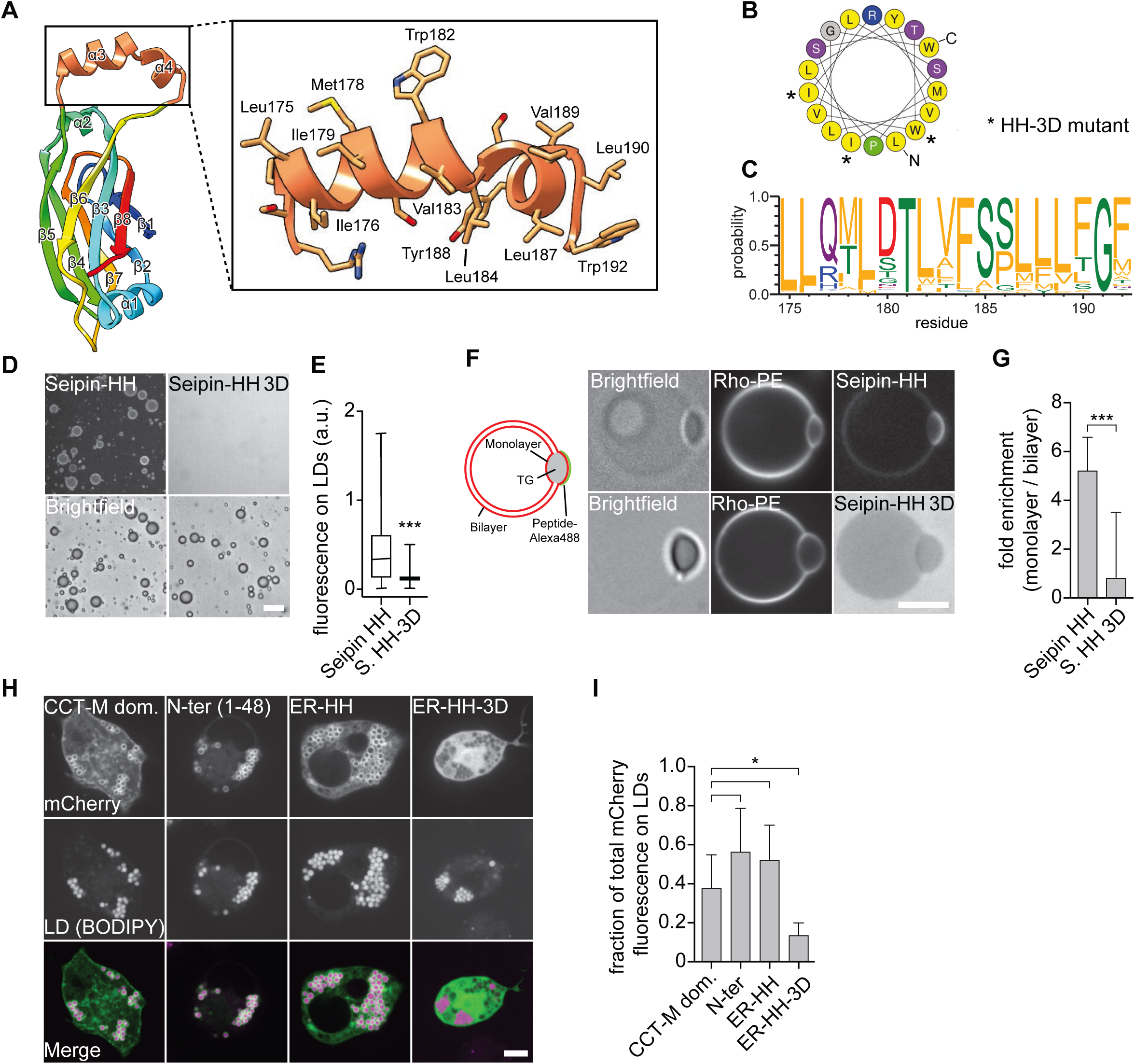
The hydrophobic helix of the seipin ER luminal domain targets to LDs. **(A)** The molecular structure of the *D. melanogaster* hydrophobic helix highlighting residues spanning 172–192 in orange. **(B)** Helical plot of residues Leu175–Trp192. Non-polar residues are shown in yellow (Gautier et al., 2008). Asterisks indicate residues mutated to Asp in the seipin-3D mutant (see below). **(C)** The helical region residue distribution for the top 200 seipin sequences (retrieved from Pfam database, corresponding to residues 175–192 of *Drosophila* seipin) shows evolutionary conservation of hydrophobicity (Crooks et al., 2004). Residues are colored according to their physicochemical properties, with hydrophobic residues in orange. **(D)** The seipin hydrophobic helix binds artificial LDs *in vitro*. An Alexa488-labeled peptide comprising residues 174–193, but not a version with the 3D mutation (replacing Ile176, Ile176 and Trp182 with Asp), binds to artificial LDs. Scale bar, 20 µm. **(E)** Quantification of fluorescent signals from >2000 artificial LDs per peptide as shown in **C**. **(F)** The seipin hydrophobic helix binds to the phospholipid monolayer *in vitro*. Seipin helix peptide, but not the mutated 3D version, preferentially binds to the phospholipid monolayer of TG lipid lenses incorporated into GUVs. A graphical representation, as well as representative confocal images show the peptide and phospholipid signals, respectively. Scale bar, 5 µm. **(G)** Quantification of enrichment on monolayer vs. bilayer of ≥18 GUVs per peptide as mean +/-standard deviation. **(H)** Binding of seipin hydrophobic helix to LDs in cells. The mCherry-tagged N-terminal amphipathic helical sequence (1–48), the luminal hydrophobic helix (174–193) of seipin, and a seipin-HH 3D mutant of the luminal helix were expressed in *Drosophila* S2 cells and analyzed by confocal imaging for LD binding. As a control, the CCT-M domain was expressed in S2 cells from the same vector. Scale bar, 5 µm. **(I)** Quantification of mCherry fluorescence on LDs vs. total signal per cell as shown in **H** as mean +/-standard deviation from ≥9 cells per construct.

### Luminal or N-terminal helical sequences and the luminal domain are required for lipid droplet formation

We next used the seipin structure to identify key features of the protein that are required for its function in cells. First, we tested the requirement for the hydrophobic helices by assaying seipin function in LD formation after expression of either the wildtype hydrophobic helix or a variant containing negatively charged residues (seipin-HH 3D) in SUM159 cells lacking seipin. After 24 h of oleate addition to induce LD formation, wild-type cells form many small LDs, while cells lacking seipin form numerous smaller LDs and some supersized LDs (>1.5 µm). As expected, re-expression of wild-type *Drosophila* seipin as an N-terminal GFP-fusion protein rescued the seipin knockout phenotype, whereas expression of GFP alone did not (Figs. 4 A and S3 D). To our surprise, expression of the 3D hydrophobic helix variant also rescued the seipin deletion phenotype of impaired LD formation (Fig. 4 A and B), indicating that this helical region was sufficient to bind LDs but is not required for LD formation. We previously found that the N-terminal sequence of seipin, likely forming an amphipathic helix, also was sufficient to bind LDs but is not required for LD formation (Fig. 3 H) (Wang et al., 2016). We therefore reasoned that these different LD-binding helices may function redundantly in LD formation. Indeed, expression of a double mutant of the N-terminus deletion and the 3D hydrophobic helix mutation was not able to rescue the LD phenotype of seipin knockout cells (Fig. 4 A and B). Similar to wildtype seipin, all constructs localized to the ER (Fig. 3 A). These findings suggest that the N-terminal and luminal helices have an important, redundant function in LD formation.

**Fig 4.**
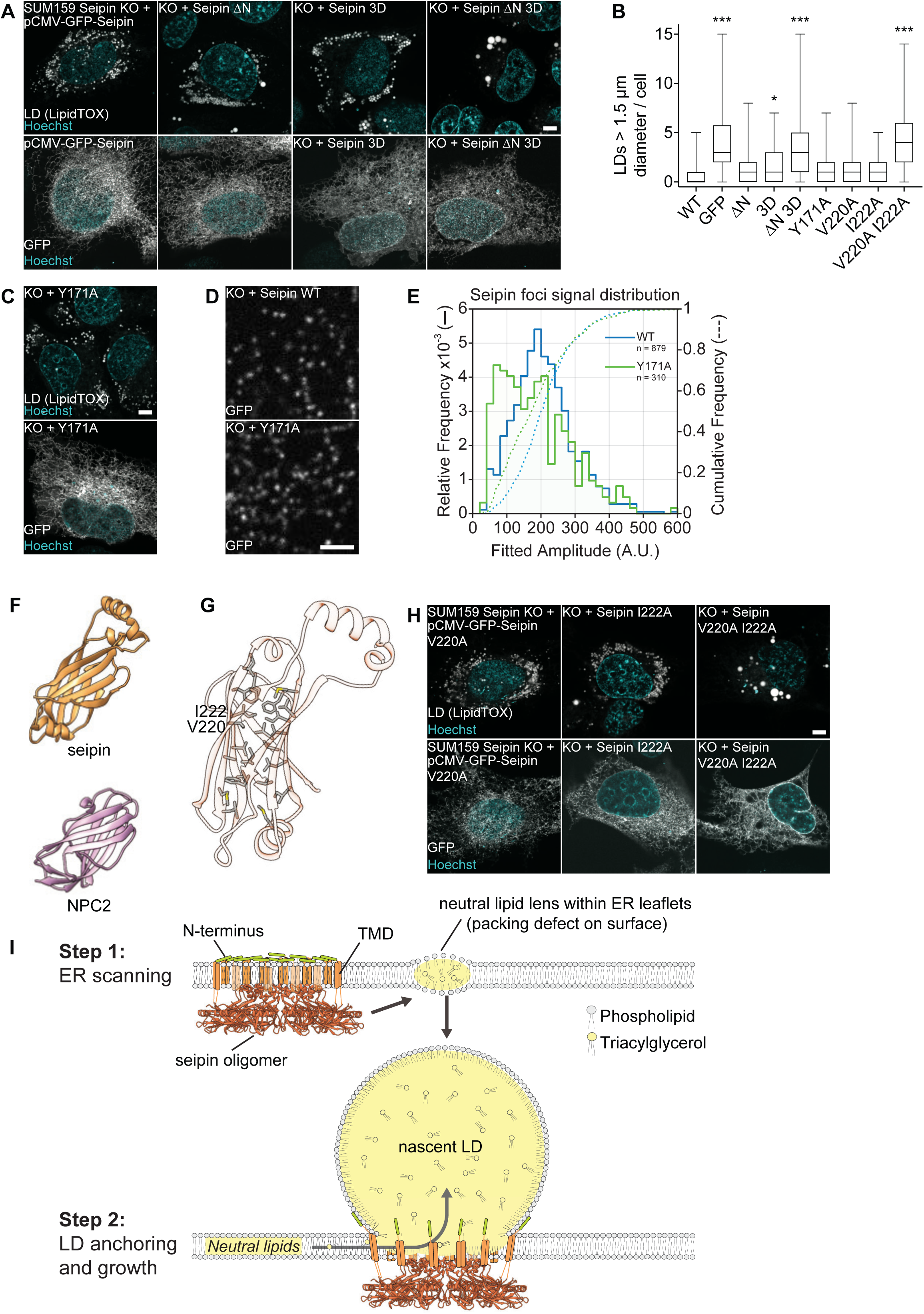
Testing key features of the seipin luminal domain in cells. **(A)** The hydrophobic and N-terminal helices are required for seipin function. SUM159 *seipin* knockout cells were transfected with the indicated seipin constructs with N-terminal GFP and analyzed for LD phenotype after 24 h of oleate treatment by LipidTOX staining (top) and localization to the ER using GFP-fluorescence (bottom). Representative images of the LD phenotype and GFP-signal are shown. Scale bar, 5 µm. **(B)** LD size of ≥47 transfected cells per construct from experiment in **A** and **C** represented as boxplots. *p<0.01; ***p<0.0001 compared to WT sample. **(C)** Seipin Y171A mutant rescues seipin deficiency. Transfection and cell treatment as in **A**. (top) LD phenotype, (bottom) localization to the ER. **(D)** Seipin wild-type and Y171A form fluorescent foci in the ER. Cells were imaged without oleate addition. To monitor pucta, low transfected cells were monitored. Scale bar 2 µm. **(E)** Seipin puncta of nine cells expressing seipin WT and 17 cells expressing seipin Y171A as in **D** were tracked and quantified over time. The comparative foci signal distributions are shown for seipin WT (blue lines) and Y171A mutant (green lines). **(F)** Structural comparison of seipin (orange) and the cholesterol binding protein Niemann-Pick type C2 (NPC2, pink; PDB accession ID: 2HKA) with a mean RMSD of 4.3Å over 106 residues is indicated. **(G)** Hydrophobic residues form a putative pocket in the luminal domain of seipin. Residues mutated in Figures 4B and 4H are indicated. **(H)** Analysis of dmSeipin luminal domain constructs as in A. **(I)** Model for the molecular function of seipin during LD formation from the ER. See text for details.

Next, we investigated whether interactions of the luminal domains are required for seipin function in LD formation by re-expressing various interface mutants in seipin-deletion cells. Each of the interface-mutant seipin proteins was able to rescue the LD phenotype of seipin-deletion cells, including those that destabilized the oligomer in *in vitro* analyses (Figs. 4 B, C, and S3 D-F). Even the Y171A mutant, which was most compromised in oligomerization *in vitro*, rescued the LD phenotype and maintained the ability to form foci in the ER, likely indicating the assembly of oligomers or higher order complexes (Fig. 4 D). Tracking of seipin foci of wild-type and the Y171A mutant over time revealed that the Y171A mutant was present in foci with similar intensity as wild-type seipin, but also in another population with substantially less GFP intensity (Fig. 4 E), suggesting seipin foci with fewer seipin molecules. Taken together, the results of expressing interface mutants suggest that interactions of the luminal domains likely contribute, but are not strictly required for seipin oligomerization *in vivo*.

Based on the seipin structure, we also tested the requirement of the highly conserved luminal domain for seipin function. This domain has a β-sandwich fold with similarity to other lipid-binding domains, *e.g.*, that found in the lysosomal lipid carrier protein NPC2 (Fig. 4 F). To test whether this domain is required for seipin function in LD formation we mutated V220 and I222 (to alanines) at the center of a hydrophobic cavity within the β-sandwich domain (Fig. 4 G and H). Although expression of either single mutant was able to rescue the seipin-deletion phenotype, expression of the double V220A and I222A mutant failed to complement the phenotype of seipin knockout cells, highlighting the importance of the luminal domain to seipin function.

## DISCUSSION

In this study, we report a molecular structure for *D. melanogaster* seipin and test features of this structure both *in vitro* and in cells. Our data indicate that *D. melanogaster* seipin forms a large, ring-shaped dodecamer with N-and C-terminal segments oriented towards the cytoplasm, 24 transmembrane domains, and an assembly of folded domains that are localized in the ER lumen. The structure reveals that the macromolecular complex positions 24 hydrophobic helices of both the luminal and N-terminal domains on either side of the ER membrane, possibly to detect and bind neutral lipid lenses forming in the ER bilayer. The dimensions of this complex are consistent with a barrel-shaped ring that connects the ER membrane with nascent LDs during their formation. The molecular structure of seipin argues strongly that seipin performs a structural, and possibly a lipid transfer, role in organizing LD formation, and suggest that other effects of seipin deficiency, such as changes in ER calcium homeostasis found with seipin deficiency, are indirect.

Many avenues of evidence indicate that the seipin luminal domain is crucial for its function. This domain is highly evolutionarily conserved and is the location of numerous lipodystrophy mutations (Magre et al., 2001a), and we demonstrate that mutating the *Drosophila* seipin luminal domain impairs its function in LD biogenesis. Our cryo-EM structure of the *D. melanogaster* luminal domain now defines structural features of this domain that begin to shed light on how seipin functions. Most clearly, our structure suggests that seipin oligomers position multiple hydrophobic helices on either side of the ER membrane, possibly to detect phospholipid packing defects due to forming neutral lipid lenses. This is consistent with previous studies that indicate that seipin foci and nascent LD foci are initially separate but can interact and subsequently co-localize during LD formation (Grippa et al., 2015; Salo et al., 2016; Wang et al., 2016). An advantage of an oligomeric structure of multiple LD-binding helices is that it increases the ability of such helices to detect lipid lenses through increased avidity, a property that we showed is relevant to the binding of amphipathic helices to LD surfaces (Prevost et al., 2018). Whether a dodecameric oligomer is strictly required for this function is in question. Data from other species suggest the number of monomers can vary between species (unpublished observations), suggesting there is flexibility in the numbers of seipin molecules in a macromolecular structure. Also, we found that mutations at the luminal interface weakened oligomerization *in vitro* but could still rescue the seipin-deficiency phenotype in cells. However, the latter finding may indicate that these mutations, while sufficient to weaken oligomerization when analyzed biochemically, do not sufficiently reduce the affinity between monomers to abolish oligomerization in the ER membrane (as shown in Fig. 4 D). Conceivably, other regions of the protein, such as interactions between transmembrane domains, may contribute to oligomerization in cells and may be essential for seipin function.

Our structure also suggests that the luminal domain may have functions that are independent from providing an anchor for the complex on the luminal side of the ER membrane. An intriguing possibility is that the luminal domain functions to mediate lipid transfer to growing LDs. The β-sandwich fold of the luminal domain is structurally similar to NPC2, which binds and solubilizes cholesterol in the lumen of the lysosome to deliver it to membrane-embedded NPC1 for export from this organelle to the cytoplasm (Wang et al., 2010). Other distantly related NPC2-type proteins (e.g., in *Camponotus japonicus*) have a similar flexible β-sheet sandwich structure that allows binding of semiochemicals, including fatty acids, used for chemical communication (Ishida et al., 2014). Our structure, and alignments with sequences from other species, suggest that like NPC2, seipin may have a binding pocket of sufficient size for accommodating hydrophobic molecules (Fig. 4 F). Perhaps the NPC2-like luminal domain of seipin participates in transferring lipids from the ER luminal leaflet to the nascent LDs to maintain the proper balance or composition of phospholipids or neutral lipids. The identity of a lipid that binds to seipin is currently unknown and under investigation.

Based on the seipin structure, we suggest a new model for LD formation (shown in Fig. 4 I). In this model, seipin forms an oligomeric complex in the ER that moves throughout the reticular network in the absence of LDs. When neutral lipids are synthesized, they are initially dispersed within the ER bilayer. However, once the TG concentration in the ER exceeds a critical concentration, lipid lenses form and locally disrupt phospholipid bilayer packing of the ER membrane, resulting in localized surface defects, similar to those on mature LDs (Prevost et al., 2018). Seipin complexes may recognize the phospholipid packing defects at these lenses by binding via its many amphipathic and hydrophobic helices located at the cytoplasmic N-terminus and in the ER lumen, respectively, and subsequently a seipin oligomer becomes localized to a neutral lipid lens (Wang et al., 2016). As nascent LDs grow toward the cytosol, seipin may function as a “molecular washer”, effectively anchoring the nascent LD to the ER (via N-terminal helix binding) and allowing for maintenance of the ER-LD connection. This could stabilize ER-LD connections, enabling LD growth and preventing nascent LDs from premature severing, as found with seipin deficiency (Grippa et al., 2015; Salo et al., 2016; Wang et al., 2016). In this model, oligomerization could also serve to restrict the diameter of the neck of the budding LDs.

In summary, we have determined a structure for one of the key proteins that is crucial for LD formation, providing an initial biochemical and structural glimpse into this fascinating process. The structure reveals an oligomeric complex, and we identify key features of this complex that participate in LD formation, suggesting a structural and/or lipid-transfer function for seipin that can be further tested. Our structural model provides a new framework for the further molecular dissection of the LD formation pathway.

## Acknowledgments

We thank Drs. Zhiheng Yu and Hui-Ting Chou from cryo-EM facility at HHMI Janelia Research Campus and Drs. Chen Xu and Kangkang Song at University of Massachusetts for cryo-EM data collection, and Dr. Srigokul Upadhyayula for help with image data analyses. We also thank members of the Liao and Farese & Walther laboratories for discussions, Tom Rapoport for a critical reading of the manuscript, and Gary Howard for editorial assistance. This work was supported by 1R01GM123089 (to F.D.), 1R01GM124348-01 (to R.V.F) and 1R01GM097194 (to T.C.W). T.C.W is an investigator of the Howard Hughes Medical Institute. X.S was supported by the American Heart Association postdoctoral fellowship (18POST34030308). H.A was supported by a DFG research fellowship (AR1164/1-1).

The authors declare no conflict of interest.

## Author contributionss

X. Sui, H. Arlt, R. Farese Jr., M. Liao and T. Walther conceived the project. X. Sui, H. Arlt, R. Farese Jr., M. Liao and T. Walther designed experiments, and X. Sui, H. Arlt performed the experiments. F. DiMaio helped with protein *de novo* structure model building. K. Brook and D. Marks performed the evolutionary analysis of protein structure. Z. Lai performed mass spectrometry analyses of protein samples. T. Walther, R. Farese Jr., X. Sui and H. Arlt wrote the manuscript. All authors analyzed and discussed the results and contributed to the manuscript.

## Material and Methods

### Seipin expression and purification

The sequence of *Drosophila melanogaster* seipin (FlyBase ID: FBpp0070426) was codon optimized for bacterial expression and synthesized and cloned into the pET28a(+) expression vector with the enzyme restriction sites of NcoI and NotI to produce C-terminally 6×His tagged seipin. The integrity of the plasmid was confirmed by sequencing. The plasmid was transformed into the BL21(DE3) *Escherichia coli* strain (New England Biolabs) for protein expression. One-liter cultures of LB medium containing 50 µg of kanamycin/mL were grown at 37°C to an OD_600_ of 1.5– 1.7, and the culture was transferred into to 4°C for 15∼20 min. Protein expression was induced by adding isopropyl β-D-1-thiogalactopyranoside (IPTG) (Roche) to a final concentration of 0.5 mM, and an additional 1 mL of kanamycin at 50 mg/mL into 1 L culture. After overnight growth (∼ 14 h) at 190 rpm, 16°C, cells were harvested by centrifugation, suspended in TMSG buffer (50 mM Tris-HCl, pH 8.0, 5 mM MgCl_2_, 400 mM NaCl and 10% v/v glycerol), and either stored at −80 °C or used immediately. Typically, a cell pellet from 2 L cultures was resuspended in 40 mL of TMSG buffer.

All purification procedures were performed at 4°C. For each 50-mL cell suspension, 1 tablet of protease inhibitor cocktail (Roche) was added, and the cells were lysed by sonication. The lysate was cleared by centrifugation at 11,594 g for 30 min. The membrane-containing supernatant was then centrifuged at 185,511 g for 1 h. The membrane pellet was collected and homogenized with a Dounce homogenizer in equilibration buffer (50 mM Tris-HCl, pH 8.0, 400 mM NaCl, 5 mM MgCl_2_, 5% v/v glycerol and 50 mM imidazole). Protein was extracted by adding n-dodecyl-β-D-maltopyranoside (DDM) to a 1% w/v final concentration with gentle rocking for 1 h. The insoluble fraction was removed by centrifugation at 184,000 g for 30 min. The recombinant protein in the supernatant was affinity purified using a Ni-NTA affinity resin (Qiagen). Briefly, for DDM-solubilized membrane suspension from 6 L cultures, 1 mL of pre-washed Ni-NTA resin by TMSG buffer was added into the suspension. After gentle stirring for 1 h, the resin was collected and washed by 10 bed volume of wash buffer (50 mM Tris-HCl, pH 8.0, 400 mM NaCl, 5 mM MgCl_2_, 5% v/v glycerol, 50 mM imidazole and 0.1% w/v DDM) containing 5 mM ATP. The protein was then eluted by 10 C.V. of elution buffer (50 mM Tris-HCl, pH 8.0, 400 mM NaCl, 5 mM MgCl_2_, 5% v/v glycerol, 500 mM imidazole and 0.1% w/v DDM). The eluted protein was collected and concentrated to 500 µL in a 100-kDa cutoff Amicon protein concentrator (Millipore) and loaded onto a Superose 6 10/300 GL size-exclusion column (GE Health Sciences) equilibrated with gel-filtration buffer (50 mM Tris-HCl, pH 8.0, 400 mM NaCl, 5 mM MgCl_2_ and 0.05% w/v DDM or 0.05% digitonin). The peak fractions containing seipin were pooled, flash-frozen and stored in −80 °C or placed on ice for immediate use.

### EM sample preparation and data acquisition

Negatively stained specimen were prepared by an established protocol with minor modifications (Booth et al., 2011). Specifically, 2.5 µL of purified seipin in DDM or digitonin at 0.2–0.3 mg/mL were applied to a glow-discharged copper EM grids covered with a thin layer of continuous carbon film, and the grids were stained with 1.5% (w/v) uranyl formate for 30 sec. These grids were imaged on a Tecnai T12 microscope (Thermo Fisher Scientific), operated at 120 kV and equipped with a 4k × 4k CCD camera (UltraScan 4000, Gatan). A nominal magnification of ×52,000, corresponding to a pixel size of 2.13 Å on the specimen, and a defocus of ∼1.5 µm were used to record the images.

For cryo-EM, 2.5–3.5 µL of purified seipin was applied to Quantifoil holey carbon grids (Cu R1.2/1.3, 400 mesh) glow-discharged for 30 s. Our initial trails show that very few particles appeared in the vitreous ice even with a high protein concentration of ∼ 5 mg/mL. Attempts to further increase protein concentration led to severe protein aggregation as revealed by cryo-EM analysis. To overcome this problem, the grids were overlaid with graphene oxide, according to a published protocol (Bokori-Brown et al., 2016)

(https://figshare.com/articles/Graphene_Oxide_Grid_Preparation/3178669). This treatment substantially increased the number of particles embedded in vitreous ice. Optimal particle distribution was obtained with a protein concentration of 0.5–1.5 mg/ml. After applying the protein, the grids were blotted with a Whatman filter paper (grade 595) for 3 s with 90% humidity and plunge-frozen in liquid ethane cooled by liquid nitrogen, using a Vitrobot (Thermo Fisher Scientific) or Cryoplunge 3 System (Gatan). Cryo-EM data were collected on a Titan Krios electron microscope (Thermo Fisher Scientific) at the Howard Hughes Medical Institute cryo-EM facility at the Janelia Research Campus.

Image were recorded using SerialEM (Mastronarde, 2005) and a K2 Summit direct electron detector (Gatan) in super-resolution counting mode. Refer to Table S1 for more information about data collection.

### EM data processing

For negative-stain EM data, the images were binned over 2 × 2 pixels, yielding a pixel size of 2.13Å, for further processing using Simplified Application Managing Utilities for EM Labs (SAMUEL) scripts (Liao et al., 2014). For cryo-EM data, drift correction was performed using MotionCor2 (Zheng et al., 2017), and images were binned 2 × 2 by Fourier cropping to a pixel size of 2.62 Å. The defocus values were determined using CTFFIND4 (Rohou and Grigorieff, 2015) and motion-corrected sums without dose-weighting. Motion-corrected sums with dose-weighting were used for all other image processing. Particle picking was performed using a semi-automated procedure (Ru et al., 2015). 2D classification of selected particle images were carried out by ‘samclasscas.py’, which uses SPIDER operations to run 10 cycles of correspondence analysis, *K*-means classification and multi-reference alignment, or RELION 2D classification (Scheres, 2012a; Scheres, 2012b). Initial 3D models were generated with 2D averages using SPIDER 3D projection matching refinement (samrefine.py), starting from a cylindrical density that mimics the general shape and size of seipin. 3D classification and refinement were carried out using “relion_refine_mpi” in RELION. One round of 3D classification without applying symmetry was performed on the total 270,716 particles to remove bad particles and to select particles with homogenous signal for the middle stacks. Subsequently, particles from classes #3 and #5 were combined for one round of 2D classification, followed by 3D refinement with D12 symmetry applied. The resulting model showed high signal-to-noise ratio in the middle stack region, whereas the distal region exhibited weak density. The next round of 3D classification focused on the middle stack region (red mesh) and yielded a total of six classes. Among them, particles belong to class #4 were used to produce the final seipin cryo-EM map with an overall resolution of 4Å. All refinements followed the gold-standard procedure, in which two half datasets are refined independently. The overall resolutions were estimated based on the gold-standard Fourier shell correlation (FSC) = 0.143 criterion. Local resolution variation of cryo-EM maps was calculated using “relion_postprocess_mpi” with “--locres” option. The amplitude information of the final maps was corrected by applying a negative B-factor using “relion_postprocessing” with “--auto_bfac” option. The number of particles in each dataset and other details related to data processing are summarized in Table S1.

### Model building and refinement

The seipin monomer and dimer maps were segmented and extracted in UCSF Chimera by using the integrated program Segger (Pintilie et al., 2010). The seipin monomer map was used to build the seipin model. *Ab initio* model building was carried out in COOT (Emsley and Cowtan, 2004). The regions with high resolution in the monomer map, including the first two layers of all four layers of seipin, which corresponding to residues from Ala125 to Glu210, could be manually built with confidence. For the low-resolution regions, the Rosetta package was used build the whole seipin model (see below). After building the model, the monomer structure was docked into dimer map and the dimer model was manually adjusted, refined in Phenix real-space refinement package (Adams et al., 2010). The refined model was visually inspected and adjusted in COOT, and the resulting map was further put back through the real-space refinement procedure to undergo further refinement. This iterative process was repeated until the dimer model reached optimal geometric statistics, as evaluated by MorProbility (Chen et al., 2010). Finally, the seipin dodecamer structure was obtained by docking the dimer model into the full dodecamer map in UCSF Chimera (Pettersen et al., 2004).

### Model-building with Rosetta

A poly-alanine model was initially built into the density. While secondary structure elements were clearly identified, ambiguity in loop density made topology determination and, consequently, sequence registration of the model difficult. Using the poly-alanine model as an input to guide placement, we ran de novo model building with Rosetta (Wang et al., 2015). Initially, Rosetta was able to place a sequence corresponding to 72 residues: a stretch spanning residues 127–200, and this model is consistent with abovementioned manually built model within the region. Using this model as an input for another two rounds of de novo modelling, the entire C-terminus was built (residues 201–239), and several strands of the N-terminus were built (residues 93–97, 109–114).

This monomeric model was then completed using RosettaES (Frenz et al., 2017). RosettaES showed very good convergence for all missing regions except over a stretch around residues 99–103. This corresponds to a region of relatively poorly resolved density. Based on monomeric structure energy and visual inspection, one conformation of this loop was selected, and refined (Wang et al., 2015) in the context of the C12 complex with the half of entire density map. In total, 1500 refined models were generated. Inspection of the five lowest-energy models yielded low-every good convergence (< 1 Å RMSD) except over the aforementioned loop, indicating confidence in the assigned model.

## Data availability

The cryo-EM density map of seipin has been deposited in the Electron Microscopy Data Bank under accession number EMD-####. Atomic coordinates for the atomic models has been deposited in the Protein Data Bank under accession number ####.

### Fluorescent-detection size-exclusion chromatography (FSEC) assay

The seipin coding sequence was cloned into the in-house generated pFasBacMam vector with the CMV promoter for overexpression of target protein in mammalian cells where seipin was tagged by an EGFP at the N-terminus. The plasmid was transfected into HEK293F cells by polyethylenimine (PEI), as described (Goehring et al., 2014; Kawate and Gouaux, 2006) in a six-well plate format. Cells were harvested after an ∼ 48-hr transfection and washed once by PBS, and the pellet was either stored at −80 °C or used immediately. To lyse the cells, the pellet from one well of six-well plate was resuspended in 500 µL of buffer containing 50 mM Tris-HCl, pH 8.0, 400 mM NaCl, 5 mM MgCl_2_ and 1% w/v DDM and supplemented with protease inhibitor cocktail (Roche). The mixture was place on a shaker with gentle shaking in a cold room for 1 hr, and centrifuged at 17,000 g on a precooled bench-top centrifuge for 20 min at 4°C. Then the detergent solubilized supernatant was collected, injected into an HPLC system (Waters) equipped with a fluorescent detector with excitation and detecting wavelengths of 488 and 520 nm, respectively. The total 20-µL protein sample was injected into the HPLC system coupled with a reverse phase gel-filtration column (5 μm, 7.8 × 50 mm) with a pore size of 500Å (Sepax Technologies). Gel-filtration analysis was performed with the running buffer containing 50 mM Tris-HCl, pH 8.0, 400 mM NaCl, 5 mM MgCl_2_ and 0.05% w/v DDM. To generate different seipin mutants, the QuickChange Site-Directed Mutagenesis kit (Agilent) was used with the protocol provided by the manufacturer.

### Cell culture and transfection

For *Drosophila* S2 cells, transfection and oleic acid treatment were performed as described (Prevost et al., 2018) with the following modifications: the cells were treated with oleic acid overnight, starting 6∼24 hours after transfection with plasmids encoding mCherry-tagged constructs. Oleic acid (complexed to BSA at a 3:1 molar ratio) was used at a concentration of 1 mM. For experiments with SUM159 cells, the culture were grown as described (Jayson et al., 2018). For seipin rescue experiments, cells were transfected using FuGENE HD transfection reagent (Promega) 1 day before addition of 0.5 mM oleate to induce LD formation. Before imaging, cells were washed in medium containing DMEM/F12 without phenol red (Thermo Fisher) and stained with LipidTox deep red and Hoechst dyes (Thermo Fisher).

### Microscopy

Imaging experiments were performed on Nikon Eclipse Ti inverted microscopes equipped with CSU-X1 or W1 spinning-disc confocal scan heads (Yokogawa), 405-, 488-and 639-laser lines, 20x Plan Apo 0.7 NA, 60x Plan Apo 1.40 NA or 100x ApoTIRF 1.4 NA objectives (Nikon, Melville, NY), and Zyla 4.2 Plus sCMOS or iXon 897 EMCCD cameras (Andor, Belfast, UK).

### *In vitro* assays

Giant unilamelar vesicles (GUVs) and *in vitro* droplets were generated as described (Prevost et al., 2018). The phospholipid composition was (1-palmitoyl-2-oleoyl-sn-glycero-3-phosphocholine (POPC): 1,2-dioleoyl-sn-glycero-3-phosphoethanolamine (DOPE): L-a-phosphatidylinositol from liver, bovine (liver PI) 65: 27: 8. For GUV experiments, 0.1 mol % 1,2-dioleoylsn-glycero-3-phosphoethanolamine-N-(lissamine rhodamine B sulfonyl) (Rhodamine PE) was added to the phospholipid mixture. GUVs were incubated with 5% triolin in buffer (20 mM Tris, pH 7.5 100 mM NaCl, 400 mM glucose) for 10 min. All Alexa488 labeled peptides used in this study were purchased from Bio-Synthesis and dissolved in DMSO. For binding assays, 1 µM peptide was added to GUVs or LDs and incubated at least 5 min before imaging.

### Image analysis

Binding of peptides to *in vitro* generated LDs was quantified using Cellprofiler software (Carpenter et al., 2006). Fluorescence intensity on each LD (ring structure segmented by brightfield images) was quantified measuring integrated fluorescence intensity (illumination corrected and background subtracted), normalized by LD area.

GUV-binding assays were quantified manually using FIJI software (Schindelin et al., 2012). Average maximal fluorescence intensities on monolayer and bilayer areas of GUVs were quantified (background subtracted) to calculate enrichment on the monolayer.

For seipin rescue experiments, transfected cells were automatically detected in the GFP-channel, and LD size and number per cells were measured in LipidTOX-channel with a Cellprofiler workflow. For untransfected SUM159 wild-type cells, cell area was segmented using LipidTOX signal.

Localization of mCherry-tagged constructs to LDs in *Drosophila* S2 cells was quantified in Cell profiler by segmenting LDs in the BODIPY channel and measurement of background subtracted mCherry signal in this region vs. total cellular fluorescence.

For tracking of seipin foci, cells were optically sectioned using a spinning disc microscope to capture most of the thin periphery of the cell, where the ER is organized in a relatively planar network. The images were acquired continuously using 57 ms exposures for 4.9 s total using EMCCD camera. Prior to the analysis of the seipin foci intensities, the cells were cropped such that the region of interest was limited to the planar ER network. Fluorescence intensities of the diffraction-limited seipin foci were detected and quantified by fitting a 2D Gaussian function using theoretically approximated sigma values for imaging conditions (Aguet et al., 2013). The detected puncta were then tracked over time, and subsequently filtered to extract only those events that were tracked for longer than 1.7 s and that did not merge or split with other foci. Using a custom MATLAB script, the max fitted amplitude was extracted for each trajectory from datasets derived from seipin knockout cells expressing WT seipin or seipin Y171A and plotted as the relative and cumulative frequency distributions.

### Statistical analysis

Statistical significance of data from *in vitro* LD binding experiments and *in vivo* protein-localization and rescue experiments using GFP-tagged seipin constructs was determined by Kruskal-Wallis test for non-normally distributed data, followed by Dunn’s multiple comparisons test. GUV-binding experiments were analyzed by Mann-Whitney test in GraphPad Prism 7. For all analyses, P-values <0.01 were considered significant.

## Supplemental Figure Legends

**Figure S1.**
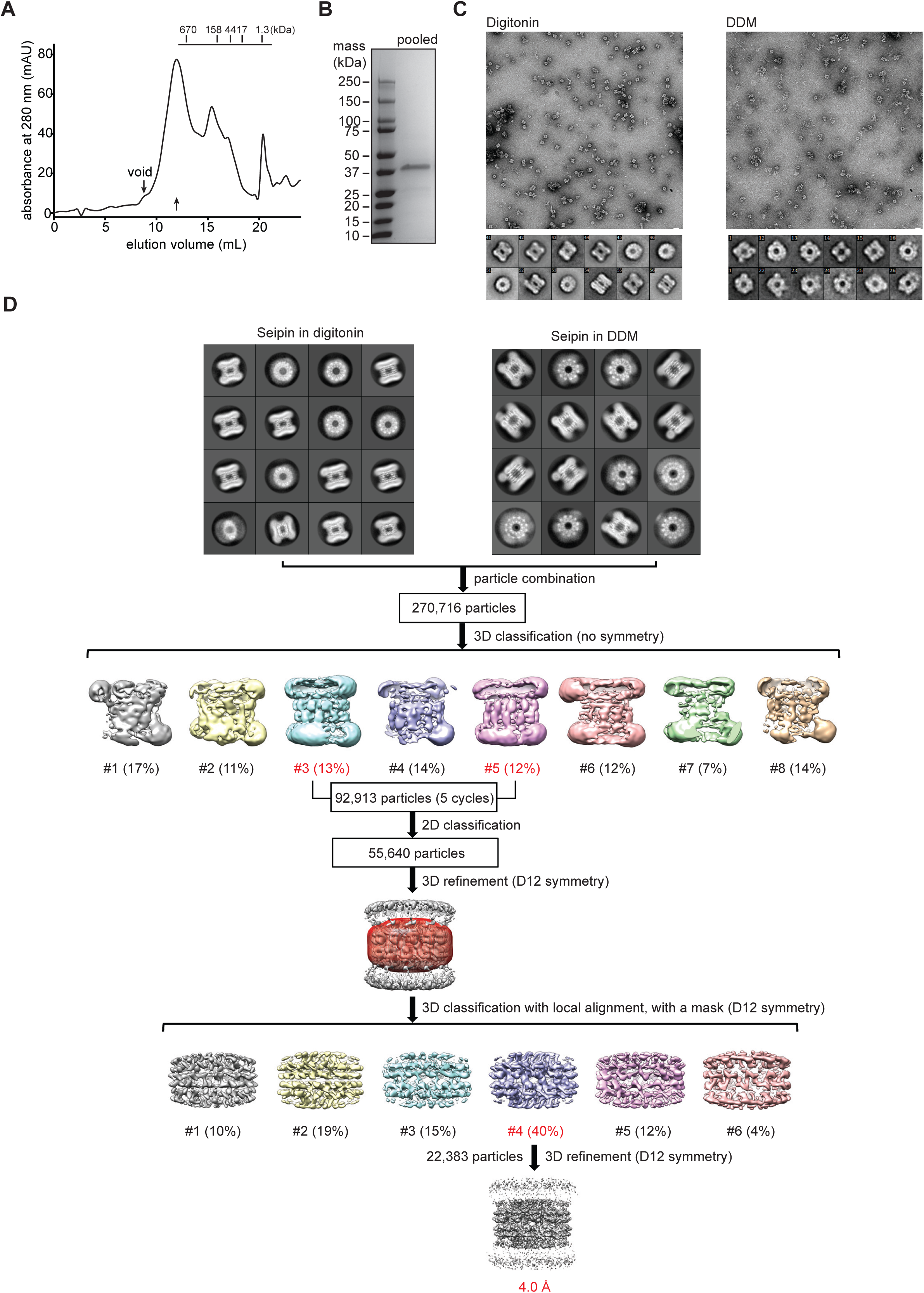
Purification, negative-stain EM analysis and cryo-EM data processing of seipin purified in digitonin and DDM. **(A)** Gel-filtration profile of seipin in digitonin as the last purification step. Only the profile in digitonin is shown, as the elution profile in DDM is similar to that in digitonin. Arrows indicate the void peak and peak containing seipin, respectively. The molecular mass label on top shows the elution peaks for protein standards. **(B)** SDS-PAGE analysis of pooled samples as indicated by arrow in panel **A**. **(C)** Representative negative-stain EM images and 2D averages of seipin (lower panel) in digitonin and DDM. Scale bar: 200 Å. **(D)** 2D average (left) of cryo-EM particle images of purified seipin in digitonin (**A**) and DDM (**B**) and single-particle EM analysis of seipin. 2D classification shows that seipin in both detergents exhibit similar overall structure. Particles collected from two detergent are combined and analyzed for subsequent cryo-EM data analysis. Workflow of image processing is illustrated. The box dimension of cryo-EM 2D average in both detergent is 393 Å. Refer to the methods section for data processing details.

**Figure S2.**
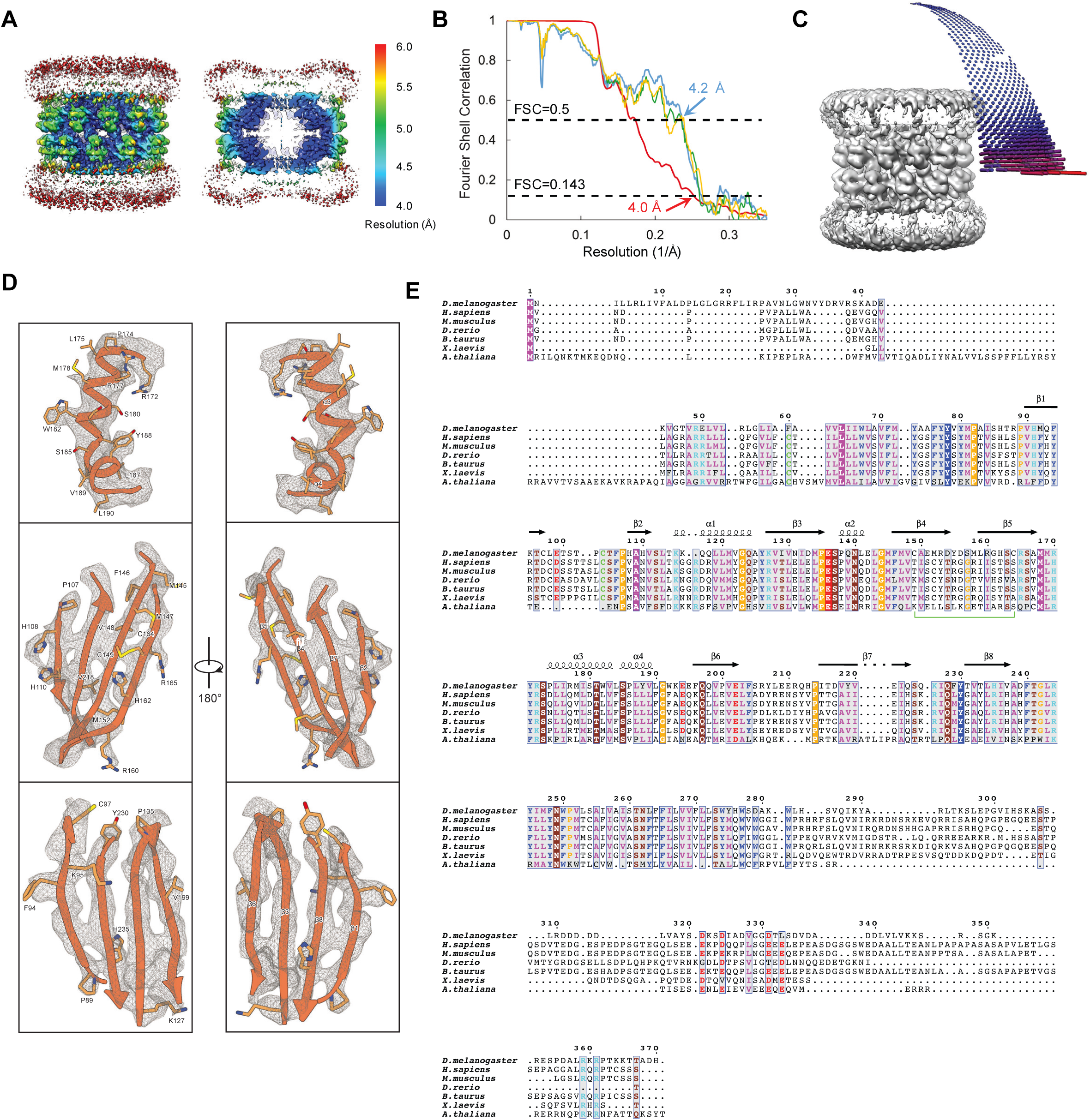
Cryo-EM 3D reconstruction of seipin. **(A)** Surface and cross-sectional views of the final cryo-EM density map filtered to 4Å. The map is colored according to its local resolution. **(B)** Fourier shell correlation (FSC) curves: gold-standard FSC between two half maps with indicated resolution at FSC=0.143 (red); FSC between atomic model and the final map with indicated resolution at FSC=0.5 (blue); FSC between half map 1 (orange) or half map 2 (green) and the atomic model refined against half map 1. **(C)** Angular distribution of the cryo-EM particles included in the final 3D reconstruction. **(D)** Selected cryo-EM densities (grey mesh) superimposed with the atomic model in different views. The densities shown here cover almost the entire atomic model of the seipin monomer. The hydrophobic helices region in the map shows the highest resolution. Lower panel shows the density matching each of the four antiparallel β-strands packed together in the seipin model. Representative residues with clear side-chain densities are labeled. The clear separation of β-strand densities demonstrates that the overall resolution of the map is consistent with the reported 4 Å. **(E).** Sequence alignment of seipin with fly seipin structure info labeled the top. The color scheme of amino acids is based on their physicochemical properties. The conserved residues among seipins from different species are boxed, and the green line indicates a pair of cysteines forming a disulfide bond in fly seipin. The structural information and residue numbering for *Drosophila* seipin are labeled on the top. Each seipin sequence was retrieved from UniProt server, and the sequence alignment was performed with T-COFFEE (Notredame et al., 2000). The final alignment figure with structural information incorporated was generated with ESPript 3.0 (Robert and Gouet, 2014).

**Figure S3.**
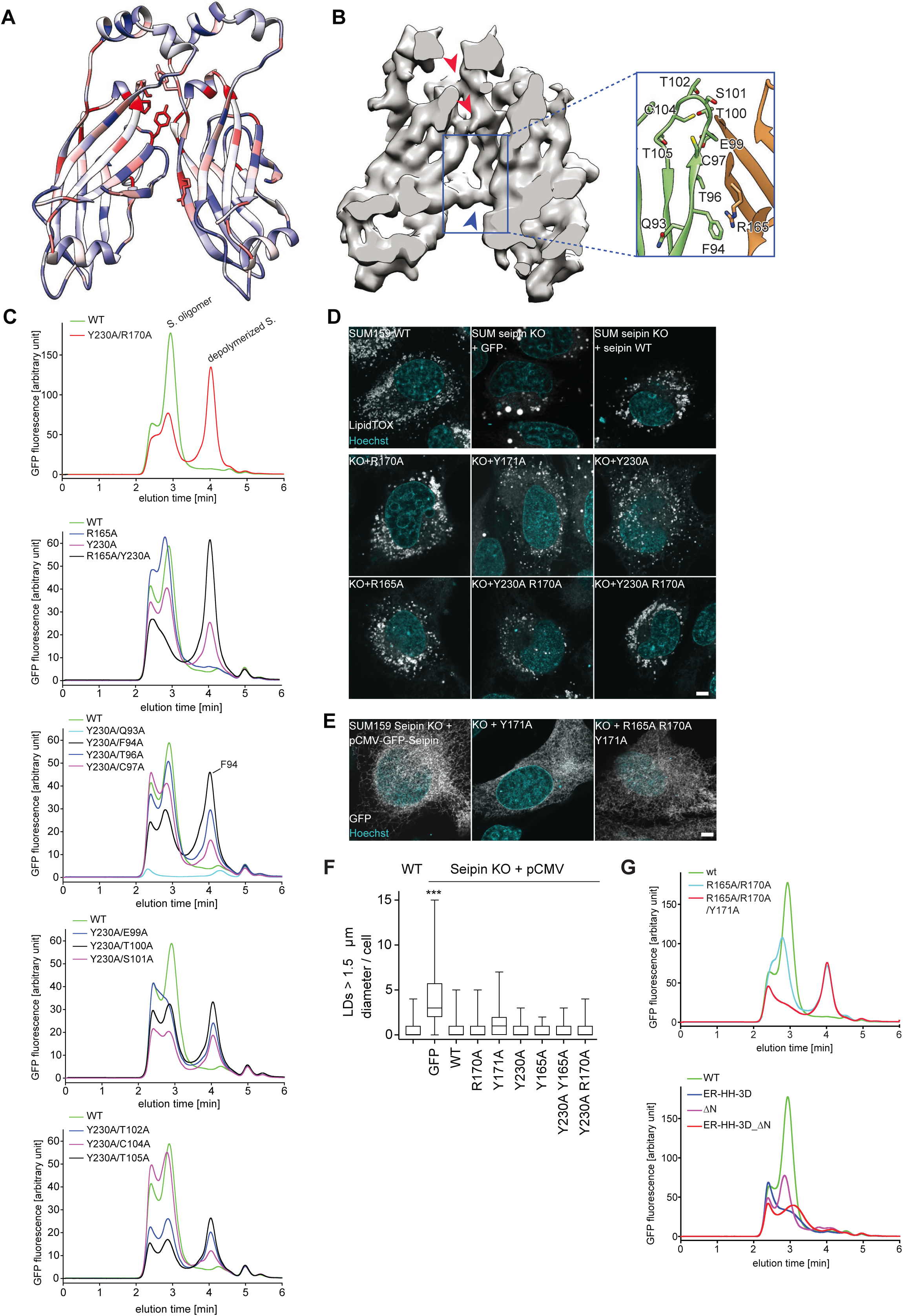
Conservation and analyses of the seipin oligomer interface and LD formation assay with selected mutant involving in seipin oligomerization. **(A)** Evolutionary conservation of seipin mapped onto the structure of a seipin dimer. Red indicates conserved residues, and blue indicates residues with low conservation, calculated from 400 seipin sequences retrieved from the Pfam database using *Drosophila* seipin to search. **(B)** Molecular model of interactions between two seipin monomers. Arrowheads indicate the three connecting regions between adjacent seipin molecules in the cryo-EM maps. Enlarged views in the boxed regions show interactions between monomers at Arg165 and Phe94. **(C)** Fluorescence-based gel-filtration analyses of seipin variants expressed in cells in identifying potential interacting residue of Arg165. Residues proximity and potentially interact with Arg165 in the neighboring seipin monomer shown in **B** are selected and mutated to Ala followed by FSEC analysis. **(D,E)** Representative images of *SEIPIN* knockout SUM159 cells expressing wild-type or oligomerization-deficient seipin mutants after 24 h of oleate treatment. LD were stained with LTOX deep red. Size bar, 5 µm. The localization of selected GFP-fused mutant are shown in **E**. **(F)** Quantification of LD sizes from experiments shown in **D**. LD-size was quantified in ≥17 cells per seipin construct. Number of LDs larger than 1.5 µm diameter per cell is shown as a boxplot representation. **(G)** FSEC analysis of selected seipin mutation in oligomerization interface, hydrophobic helices region of ER luminal domain and also lacking of cytosolic N-terminal domain. See text for details.

**Table S1.** Cryo-EM data collection, refinement and validation statistics.

|  | *D. melanogaster* seipin |
| --- | --- |
| **Data collection and processing** |  |
| Magnification | 22,500 |
| Voltage (kV) | 300 |
| Electron exposure (e<sup>-</sup>/Å<sup>2</sup>) | 58 |
| Defocus range (average) (μm) | 1.0-3.5 (2.0) |
| Pixel size (Å) | 1.31 |
| Symmetry imposed for final map | D12 |
| Number of collected movies | 4,743 (DDM)/2,608 (Digitonin) |
| Initial particle number for 3D classification (no.) | 270,716 |
| Final particle for final map (no.) | 22,383 |
| Map resolution (Å) | 4.0 |
| FSC threshold | 0.143 |
| Map resolution range (Å) | 3.9-11.6 |
| Map sharpening B-factor (Å<sup>2</sup>) | - 200 |
| **Refinement\*** |  |
| Initial model used (PDB code) | NA |
| Number of protein residues | 306 |
| Number of atoms | 2,518 |
| **Geometric deviations (r.m.s.d)\*** |  |
| Bond lengths (Å) | 0.006 |
| Bond angles (°) | 1.292 |
| **Validation\*** |  |
| MolProbity score | 1.98 |
| Clashscore | 7.95 |
| Poor rotamers (%) | 0.00 |
| **Ramachandran plot\*** |  |
| Favored (%) | 90.07 |
| Allowed (%) | 9.93 |
| Outliers (%) | 0.00 |

\* Data reported here are for the seipin dimer model

